# Impact of hypoxia induced VEGF and its signaling during caudal fin regeneration in Zebrafish

**DOI:** 10.1101/105767

**Authors:** sagayaraj.R Vivek, R. Malathi

## Abstract

Hypoxia is known to play important role during various cellular process, including regeneration. Regeneration is a complex process involving wound healing and tissue repair. We propose that hypoxia might mediate regeneration through angiogenesis involving angiogenic factors such as VEGF, VEGF-R2, NRP1a during the wound healing process. We have chosen Zebrafish model to study the role of hypoxia induced regeneration. Unlike mammals Zebrafish has the ability to regenerate. Hypoxic condition was mimicked using inorganic salt cobalt chloride to study caudal fin regeneration in adult Zebrafish. Intense blood vessel formation, with increased tail fin length experimented at various time points have been observed when adult zebrafish caudal fin partially amputated were exposed to 1% CoCl_2_. Regeneration is enhanced under hypoxia, with increased VEGF expression. To study the significance of VEGF signaling during wound healing and tissue regeneration, sunitinib well known inhibitor of VEGF receptor is used against CoCl_2_-induced caudal fin regeneration. Diminished fin length, lowering of blood vessel formation was documented using angioquant software, reduction in mRNA level of hypoxia inducible factors, VEGF and other pro-angiogenic genes such as VEGF, VEGF-R2, NRP1A, FGFR2, ANGPT1 were observed, while reduction in VEGF protein was demonstrated using western blot analysis. Genistein inhibitor of HIF-1α completely arrested regeneration, with suppression of VEGF highlighting the significance of hypoxia induced VEGF signaling during fin regeneration. Our results suggest that hypoxia through HIF-1α might lead to angiogenesis involving VEGF signaling during wound healing and this might throw light on therapeutic efficacy of cobalt chloride during regeneration.

## INTRODUCTION

Understanding the mechanism of tissue regeneration is of fundamental importance and might have therapeutic applications. Regeneration is the ability of the organism to regain lost organ/tissue and restore the same size and shape of the damaged tissue. Multiple processes are involved which includes wound healing, tissue repair, blastema formation and regenerative outgrowth that involve several signaling pathways associated with gene expression. Extensive studies on tissue repair and regeneration have shown that multiple signaling pathways are involved. Cobalt chloride, nickel chloride and desferrioxamine are known as hypoxic mimicking agents (Goldberg MA et., 1998).

Hypoxia is initiated in response to the cells having low oxygen concentration and is known to regulate various cellular process including differentiation, proliferation (Sahai A et al., 1999, Forristal CE et al., 2010, Voss MJ et al., 2010, Jogi A et al., 2003, Cassavaugh J et al., 2011) and regeneration. Cells can critically judge low oxygen level, and reported to play increased angiogenic action initiated in response to promote oxygen supply, cell survival by altering gene expression through transcription and translation (Bi et al., 2005, Koritzinsky, M et al., 2006, Wouters, B.G et al., 2005 and Susanne Jung and Johannes Kleinheinz, 2013). In response to hypoxia, Hypoxia inducible factor-1α (HIF-1α) a transcription factor is effectively known to induce angiogenesis via VEGF and its downstream targets which might be significant during wound healing and tissue regeneration. Hypoxia plays a critical role in proliferation of endothelial cells (Li W et al., 2007), fibroblasts (Mizuno S et al., 2009). Moreover exposing adult Zebrafish to CoCl_2_ is known to increase the number of endothelial cells in caudal fin (Glassford AJ et al., 2007).

Zebrafish has emerged as a powerful vertebrate genetic model to study regeneration as it can regenerate one fifth of the heart ventricles, pancreas and grow caudal fin. Zebrafish offers a major tool for biomedical research to study human diseases compared to other vertebrate models. Innate capacity to restore shape and function of amputated fin, optic nerves, spinal cord, scales, heart, (Glassford et al., 2007; Jinping Shao et al., 2011), draws the focus to select zebrafish as a model organism, to study regeneration and molecular insights during regenerative angiogenesis (Bayliss et al., 2006). Lost caudal fin tissue in zebrafish is recovered through epimorphic regeneration found similar in urodele limb like wound healing, blastema formation, restoration of tissue with differentiating cells finally regeneration (Akimenko et al., 2003). Although regenerative events is limited in humans and other mammals, molecular mechanisms seem to be conserved which mkaes zebrafish an efficient model in regenerative research.

In zebrafish many signaling pathways have been implicated during tail fin regeneration, including FGF that is known to play key role in blastema formation, MAP kinase, etc. (Arsham, A.M et al., 2003), are considered. However, hypoxia-inducible factor (HIF)- VEGF pathways found to be a prominent signaling pathway towards cellular hypoxic response. Hif-1α the major transcriptional factor activated during hypoxia could regulate angiogenesis (VEGF, FLT1, FLK1, END1, ANGPT1), erythropoiesis (EPO), glucose metabolism (Glut-1), cell survival (IGF and TGF-α), apoptosis, cancer metastasis, invasion (MMP-2, cathepsin-D) with VEGF being the target gene expressed during angiogenesis. Formation of new blood vessels is a key process of angiogenesis, with complex physiological sequence leading endothelial cell ripening, proliferation. The angiogenic homeostasis is governed by many pro angiogenic and anti-angiogenic factors. VEGF, the major angiogenic factor is found to be highly commendable in regulating angiogenesis through receptors VEGFR1/VEGFR2 (Pandya NM et al., 2006, Susanne Jung and Johannes Kleinheinz, 2013) during development (Bakkiyanathan et al., 2010), regeneration, tumor growth and metastasis (Susanne Jung and Johannes Kleinheinz, 2013) by activating endothelial growth factors, transforming growth factors (Jazwinska et al. 2007), MMPS (Bai S et al., 2005). VEGF plays crucial role by potentiating micorvascular network during regeneration, via restoring, cell shape and integrity by its own tissue renovating mechanism (Rathinasamy et al., 2014; Susanne Jung and Johannes Kleinheinz, 2013). Although many signaling molecules likes fibroblast growth factor (Kenneth D. Poss et al.,2000), notch (Angel Raya et al.,2003), BMP (A. Smith et al., 2006), Wnt/*β*-catenin (Yasuhiko Kawakami^1^ et al., 2006) studied previously, we tested if hypoxia induced regeneration involves VEGF signaling and its downstream targets promoting angiogenesis a major requisite of wound healing process.

In the present study, we demonstrate that hypoxia induced angiogenesis through VEGF signaling mutually contribute to a regenerative outgrowth. Exploring caudal fin regeneration under hypoxic condition revealed clear insight into blood vessel development and proliferation and this might enhance wound healing process. We measured the length, junction and sprouting of the blood vessel formed in dorsal, cleft and ventral region of the caudal fin using angioquant software and also obtained expression pattern of, endothelial growth factor (VEGF, ANGPT1) and its receptors (FLT1, FLK1, END1), co-receptor Nrp-1a, transforming factors (TGF-β), MMPS (MMP-9,2), signaling factors (HIF-1α, DLL4, S1RP1, HER, HEY) and pro-angiogenic genes under cobalt chloride treated condition. Therefore this scientific evidence might shed light on mechanism of VEGF signaling during wound healing process and might help to develop therapeutic strategies towards HIF- VEGF induced angiogenesis during wound healing and tissue repair.

## 2. Materials and methods

### 2.1. Chemicals and reagents

CoCl_2_, Genistein, Dimethyl sulfoxide (DMSO),sunitinib (SU5416) were purchased from Sigma Aldrich Chemicals. Pvt. Ltd. (USA). Tri Reagent (Takara), Tricaine MS-222 (MP Bio Medicals), all chemicals used is of molecular grade.

### 2.2. Zebrafish maintenance

Zebrafish (wild type) were purchased from local aquarist and maintained in internal filtered water tanks at 28°C with 14 h light and 10 h light. Fishes were fed with dry flakes (Tetra) twice a day.

### 2.3. Fin regeneration Drug administration

Six months old adult Zebrafish were anesthetized using tricane (MS-222) and their caudal fin was partially amputated using razor blade and exposed to 1% cobalt chloride. These fish were allowed to regenerate for 5 days and regenerated tissue was used for further study. Regenerated caudal fin were documented on 3,5,10 dpa. Caudal fin amputated from adult Zebrafish were exposed transdermally to 1% CoCl_2_ and treated in combination with genistein, and SU 5416 which were dissolved in DMSO at stock concentrations of 100 μM and 1 μM respectively were also allowed to regenerate after 0 – 5 dpa. 10µl of required concentration of these compounds were injected (*i.p*) respectively while control fish were treated with 10 μl of 0.1% DMSO solution.

### 2.4. Morphometric analysis

The morphology of tail fin of zebrafish exposed to 1% CoCl_2_ after amputation was documented using light microscope (Euromax) at 4X resolution. Fin length was measured from the amputation site in dorsal, cleft and ventral region of the regenerated caudal fin using image focus software (Euromax).

Blood vessels were quantified in terms of length, junction and sprouting using angioquant software (Version 1.33, MathWorks).

### 2.5. Gene expression analysis using Quantitative- PCR

RNA was isolated from Zebrafish caudal fin tissue treated with compounds (1% CoCl_2_, SU 5416, genistein) and control using Tri Reagent (Takara). Concentration, purity of RNA was determined using nano drop (Thermo Scientific) and reverse transcribed. Reverse transcription was performed for VEGF, VEGF-R2, co-receptor NRPIA and other proangiogenic factors such as ANGPT1, TIE, EGFR, FGFR, MMPS, using random primers (Promega), riboblock (Thermo Scientific), reverse transcriptase (MMuLV), reverse transcriptase buffer (MMuLV), DNTP (Invitrogen), molecular grade water (Sigma) in a PCR vial and incubated at 42 ◦C. Quantitative PCR was then performed using SYBR green (KAPPA) reagent, with the primer sequences listed (Table 1) and having β- actin as internal control.

**TABLE.1.**
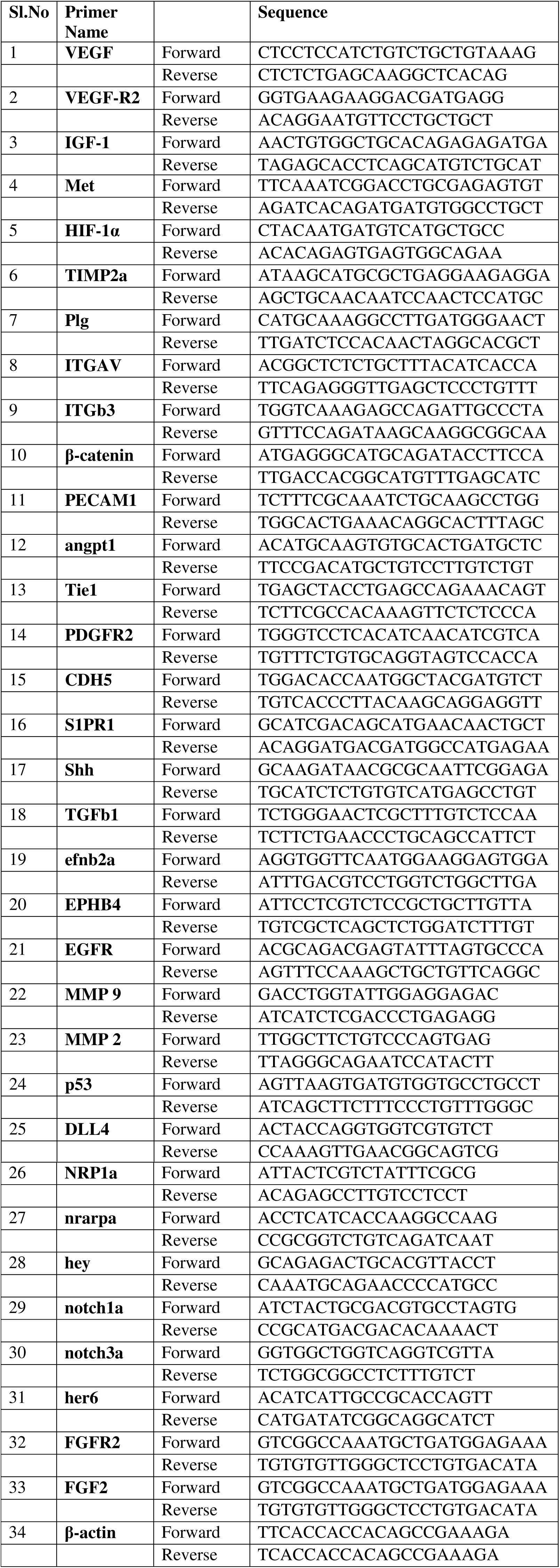
List of primer sequences of genes used in this study

### 2.5. Western blot

VEGF protein was quantified using western blot analysis. Fin tissue of adult zebrafish after regeneration in case of control receiving 0.1%DMSO, and drug treated under 1% cobalt chloride at 5 dpa was lysed and quantified using Lowry’s method.50µg protein from each groups were resolved on 10% SDS-PAGE, and subsequently transferred to PVDF membranes. Membranes were blocked with 10% skimmed milk followed by incubating with anti–VEGF antibody (R&D Systems) 1:1000 dilution at 4˚C overnight, followed by incubation with secondary horseradish peroxidase-conjugated anti - mouse secondary antibody (1:10,000 dilutions) for 1 hour at room temperature. These blots were washed with TBST and immunoreactivity was detected using chemiluminescence kit (Bio Rad).

### 2.6. Statistical analysis

Data were expressed as mean ± SEM. Statistical analyses were performed using one-way ANOVA followed by Tukey’s Multiple Comparison tests, for comparison between treated values and control values using Graph Pad Prism software. P values < 0.05, P < 0.005, P < 0.001 were considered to be statistically significant.

## 3. RESULTS

### 3.1. Hypoxia induced regeneration.

Wound healing is an important process during regeneration. To delineate whether VEGF signaling could play an important role in promoting such events, we studied regenerative cascade in relation to angiogenesis using adult zebrafish model. Zebrafish caudal fin were partially amputated at 0dpa and allowed to recover under (1%) CoCl_2_ treated condition. Tail fin length, blood vessel sprouting in regenerating fin under normoxia and hypoxia were recorded at 3, 5, 10dpa and analyzed. Zebrafish treated with 1% CoCl_2_ (1%) displayed increase in tail fin length when measured from amputation site. Caudal fin length is calculated in µm using image focus software (Euromax). Fin length significantly increased by 2035µm in dorsal, 1500µM in cleft and 1900µM in ventral regions of the caudal fin compared to the control (0.1% DMSO). Time course experiment performed at 3, 5 and 10 dpa under 1% CoCl_2_ (Fig.5) exhibits enhanced caudal fin length with significant increase in proliferating blood cells disseminating to tip of the regenerating caudal fin as revealed by microscopic (Fig.1). Fish treated upto 5 dpa was taken up for study (Fig.2). In order to find out if VEGF signaling plays an important role during regeneration, adult zebrafish caudal fin was amputated after exposing to 1% CoCl_2_ and treated with SU5416 and genistein, which are known to be inhibitors of VEGF signaling. 10µl of 25, 50, 100µM concentration of genistein was injected into abdominal region of zebrafish at 6, 18, 42, 66 and 90 hours dpa. Morphogenetic variation of regenerated caudal fin is studied from the plane of amputation. Regeneration is determined in dorsal, cleft and ventral regions of the caudal fins by measuring their length from amputated site at 5 dpa. Reduction in Caudal fin length was observed with a 100µM genistein at 310µm at dorsal, 276µm cleft and 280 µM in ventral regions compared to 25 µM genistein by 890µm, 1020 µm 860µm. At 100µM maximum inhibition rate of 84.4%, 82.2%, and 87.12% in dorsal, cleft and ventral region respectively, when compared with control receiving 0.1% DMSO (Fig.3) and 1% CoCl_2_ (Fig 6.), when fish were treated with 1µM SU 5416 an inhibitor of VEGF-R2, a reduction in fin length and number of blood vessels was observed (Fig.5). This demonstrates that sunitinib virtually completely blocked hypoxia-induced regeneration, suggesting that VEGF could be involved during wound healing process of regeneration.

**Figure.1.**
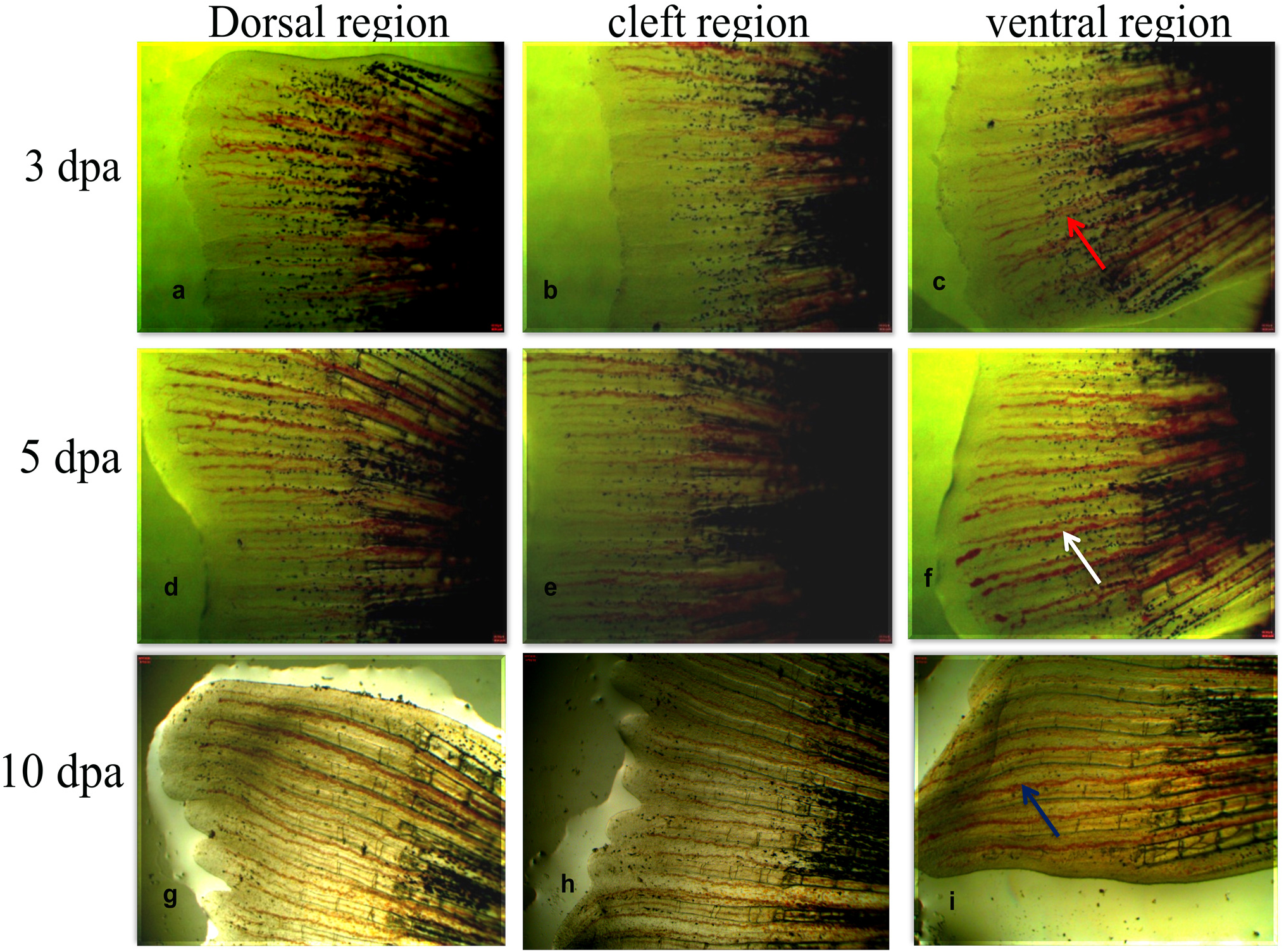
Cobalt chloride promotes blood vessel sprouting, proliferation: Representative images of the dorsal, cleft and ventral region of regenerated tail fin were acquired on 3, 5, and 10 days post amputation. Adult zebrafish caudal fin amputated transderamally exposed to 1% cobalt chloride at different time point showing increased tail fin length 2035µm in dorsal, 1500µM in cleft and 1900µM in ventral regions and blood vessel detected using light microscopy (1) Red arrow indicate blood vessel sprouting at 3dpa. (2) White arrow indicates proliferation of blood vessel at 5dpa. (3) Blue arrow indicate intense blood vessel at 10dpa.

**Figure.2.**
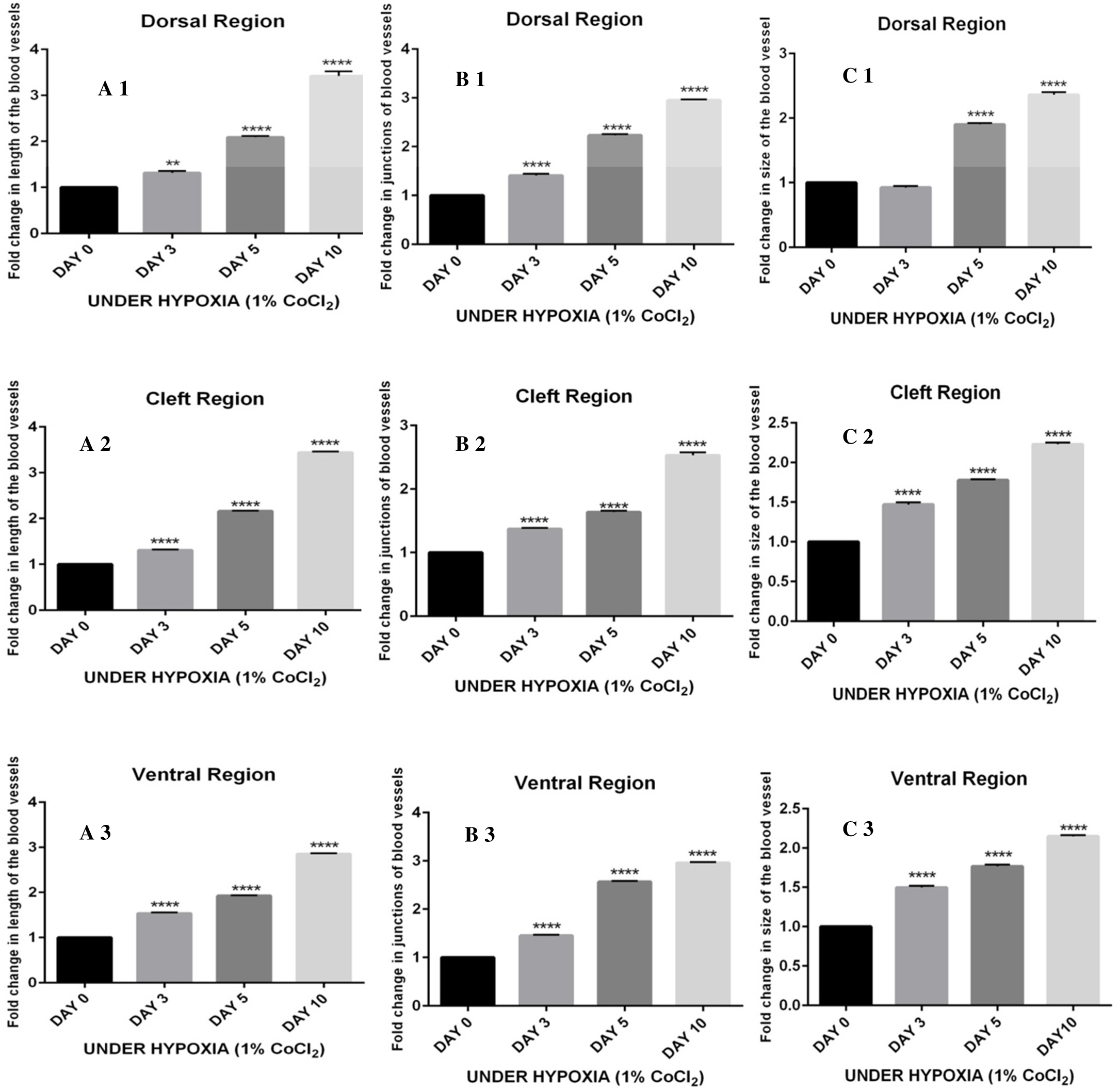
Statistical analysis of fold change in blood vessel documented using angioquant software:Increase in (A 1, 2, 3) length, (B 1,2,3) size and (C 1,2,3) Junction of blood vessel of the dorsal, cleft and ventral region in regenerated tail fin under hypoxia condition observed at 0dpa, 3dpa, 5dpa and 10dpa. At 3 dpa size of blood vessel increased by 1.4 fold in at dorsal, 1.3 fold in cleft and 1.5 fold in ventral regions. At 5dpa Size of the blood vessel showed fold increase of 1.8 times in dorsal region, 1.6 times in cleft and 1.7 in dorsal region of the caudal fin. Blood vessel length were found a fold increase of 2.3 in dorsal region 2.2 in cleft and 1.9 in ventral region, compared to normoxia, lacking signs of blood vessel development in regenerated tissue. at 10 dpa is significantly superior to control demonstrating a fold increase of 3.6 in length, 2.8 in junction and 2.6 in size (Fig.2) of the dorsal region compared to cleft region with fold increase at 3.5in length, 2.7in junction, 2.3 in size and ventral region with 2.8 in length, 2.7 in size and 2.1 in size of the blood vessel. The fold change is expressed as mean ± SEM. Statistical analyses performed using one-way ANOVA followed by Turkey’s multiple comparison tests for comparison between treatment values and control. P values < 0.05 considered to be statistically significant. ** P value <0.005, *** P value < 0.001, **** P value < 0.0001.

**Figure.3.**
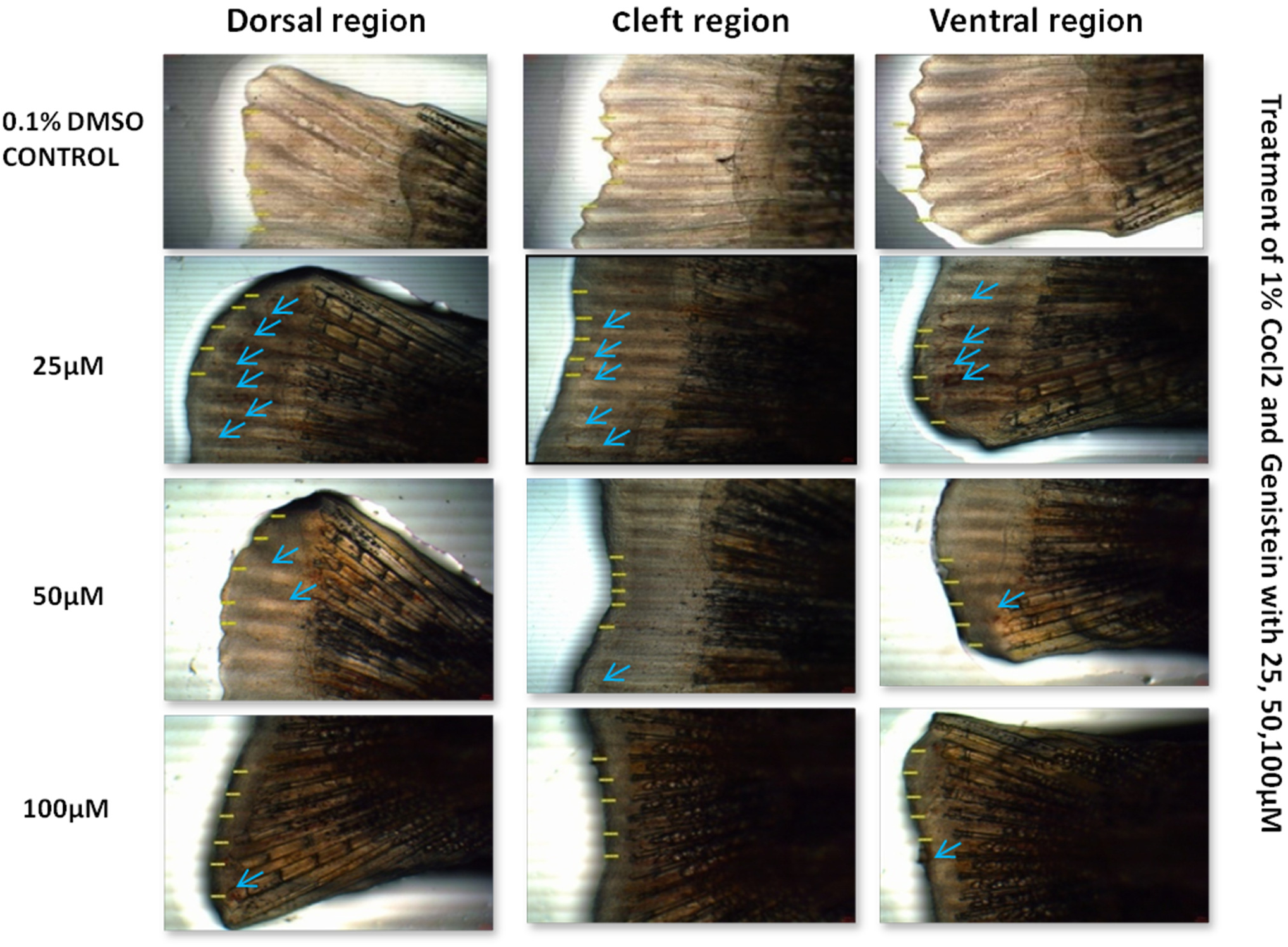
Inhibition of blood vessel sprouting and proliferation by genistein: Representative images of the dorsal, cleft and ventral region of regenerated tail fin at 5dpa under hypoxia (1% CoCl_2_) combined with varying concentration of Genistein treatment. Red arrows indicate plane of amputation and yellow arrow indicate area of regeneration. (A). INHIBITION OF CAUDAL FIN LENGTH: Reduction in caudal fin length is observed with increasing concentration of genistein at 25, 50,100 μm under 1% CoCl_2_ treatment (B). Adult zebrafish amputated caudal fin treated with or without 25, 50, 100 µm genistein. Blue arrows indicate blood vessel sprouting in the regenerated caudal fin.

**Figure.4.**
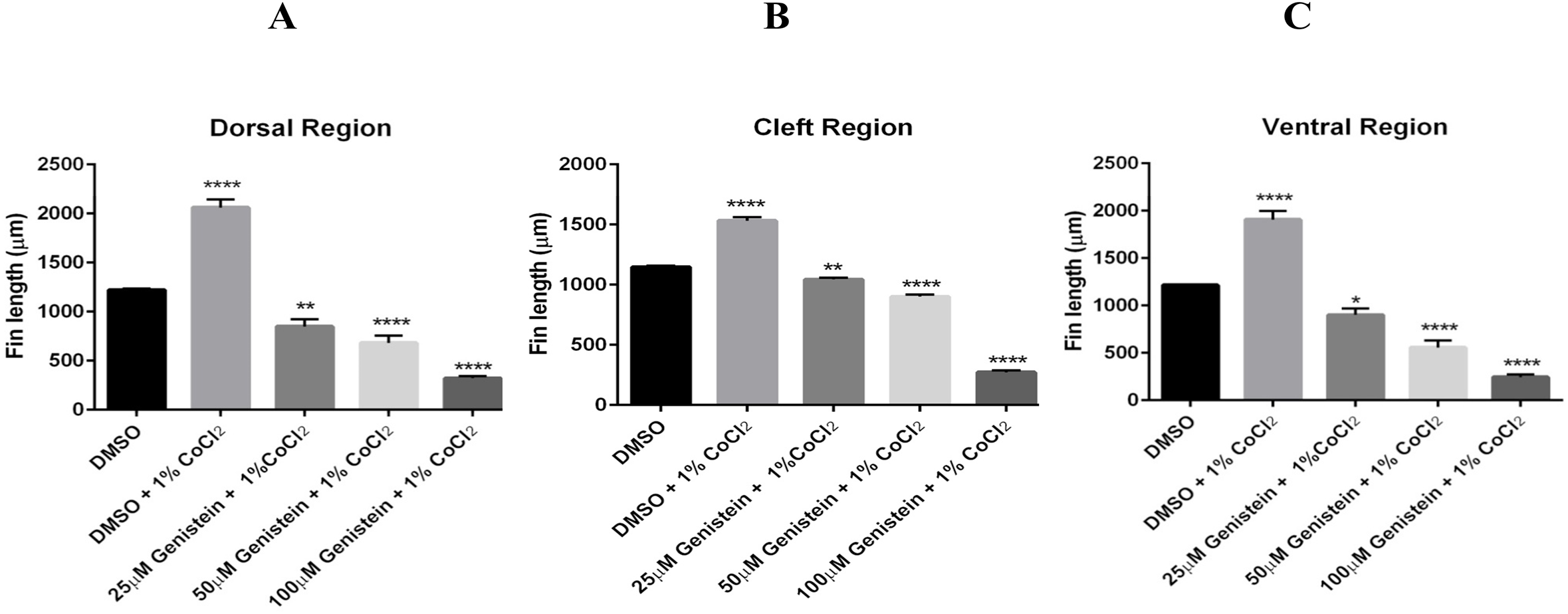
Fin length significantly increased by 2035μ m in dorsal, 1500µM in cleft and 1900µM in ventral regions of the caudal fin compared to the control (0.1% DMSO) showing 1290µm, 1090µm, and 1315μ m. On 5dpa decrease in caudal fin length is observed under 1% CoCl_2_ with increasing genistein concentration. At 25μM, μM, 100μ M fin length significantly decreased by 890μm, 775μm, 310μm at dorsal region compared to cleft region showing 1020 μm, 910μm, 280 µM and dorsal region at 860μm, 640 μm, 276μ m in ventral regions of the caudal fin. The data (μm) is expressed as mean ± SEM. Statistical analyses performed using one-way ANOVA followed by Turkey’s multiple comparison tests for comparison between treatment values and control. P values < 0.05 considered to be statistically significant** P value <0.005, *** P value < 0.001.

**Figure.5.**
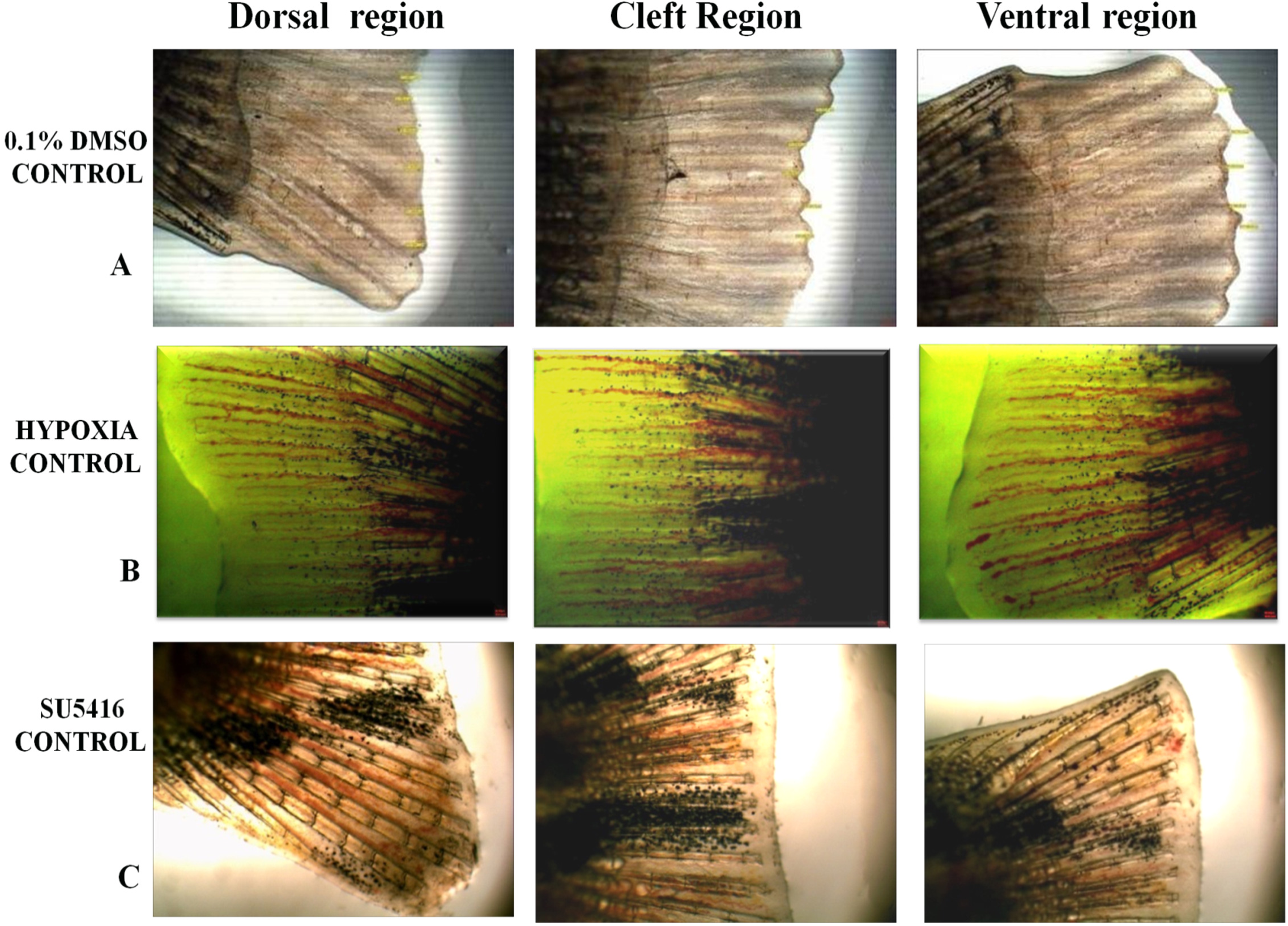
Sunitinib inhibits cobalt chloride induced blood vessel sprouting, proliferation: Adultzebrafish caudal fin partially amputated were placed in hypoxia water containing 1% cobalt chloride, followed by treatment with 10 µl of 1µm SU 5416 at 6,18,36,66,90hpa. Representative images of the dorsal, cleft and ventral region of regenerated tail fin at 5dpa under the following condition (A),Control receiving 0.1% DMSO, (B) hypoxia control (1% CoCl_2_), (C) 1µM SU 5416. Increased blood vessel sprouting and tail fin length was detected compared with control receiving 0.1% DMSO and hypoxic control at 5dpa but reduced in the presence of SU5416.

**Figure.6.**
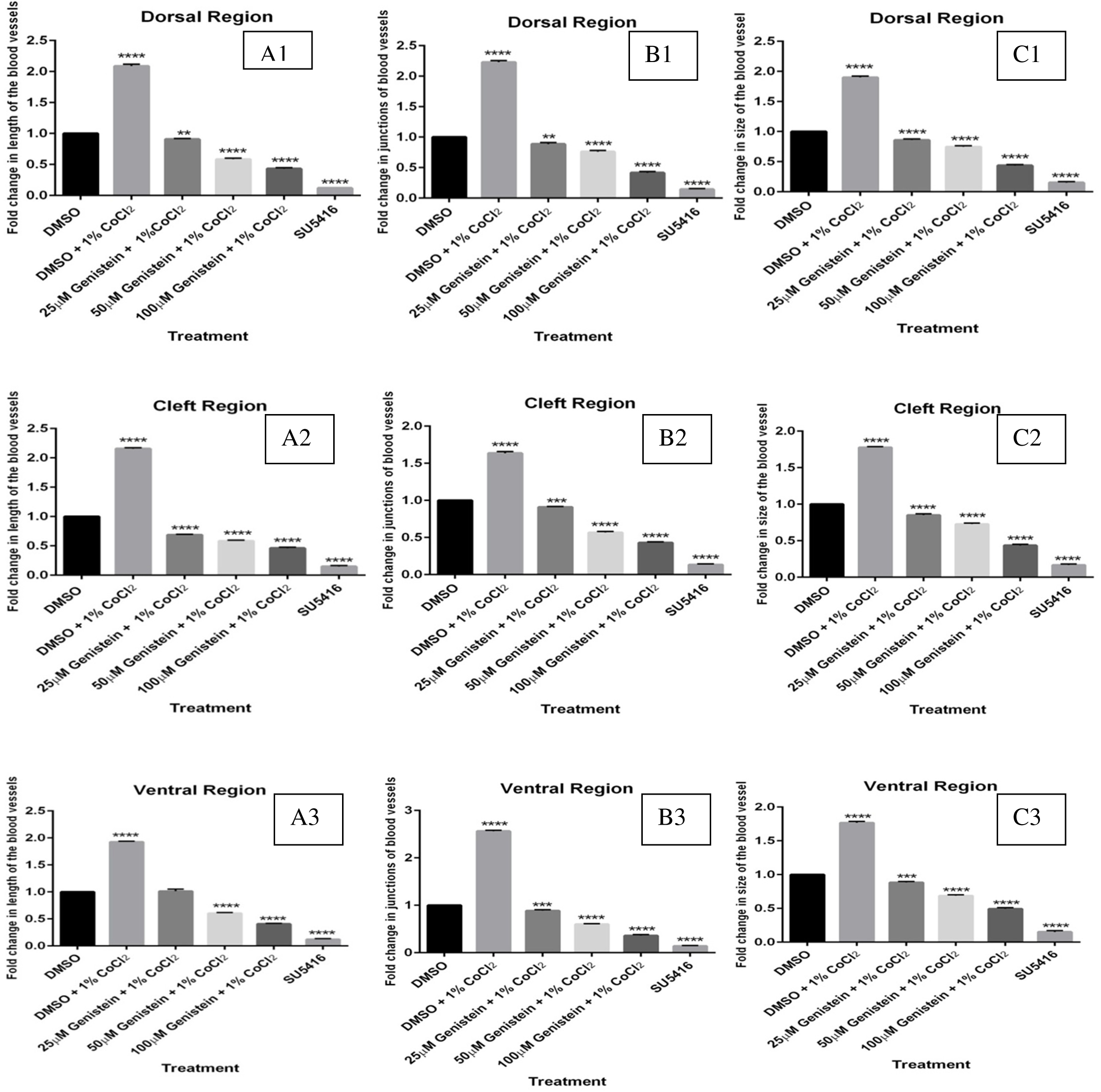
Statistical analysis of fold change in blood vessel: Decreased in (A) length, (B) size and (C) junction of blood vessel of the dorsal, cleft and ventral region in regenerated tail fin under hypoxia condition is observed at 5dpa with increasing concentration of Genistein. Blood microvasculature was significantly reduced by genistein treatment at 100µM expressing only fold change of 0.3 in length and 0.4 in size of the blood vessel at dorsal region, compared to 0.4 and 0.6 times at cleft region, and 0.34 and 0.52 in ventral region respectively, compared to more than 2 fold increase under 1% CoCl_2_ condition. Furthermore results were compared with fish treated with 1µM SU5416 measuring length of blood vessel at 0.1 fold changes in dorsal, 0.15 at cleft and 0.1 in ventral regions. Fold change in size of blood vessel was observed at 0.14 in doral, 0.2 at cleft and 0.16 at ventral regions were found significantly inhibited compared to control. The fold change is expressed as mean ± SEM. Statistical analyses were performed using one-way ANOVA followed by Turkey’s multiple comparison tests for comparison between treatment values and control. P values < 0.05 considered to be statistically significant. ** P value <0.005, *** P value < 0.001, **** P value < 0.0001

Blood vessels were quantified using angioquant software. An increase in number of blood vessels were recorded in 1% CoCl_2_ treated groups when compared to control (Fig.2). This was determined in terms of length, junction and sprouting of vessels formed in the dorsal, cleft and ventral regions of the caudal fin. At 3 dpa a fold increase of 1.4 in size of blood vessel at dorsal, 1.3 in cleft and 1.5 in ventral regions found neighboring to wounded site. At 5dpa blood vessel length were detected at a fold increase of 2.3 in dorsal region 2.2 in cleft and 1.9 in ventral region, compared to under normoxia, which shows reduction in blood vessel development in regenerated tissue. (Fig.5) and size of the blood vessel showed fold increase of 1.8 times in dorsal region, 1.6 times in cleft and 1.7 in dorsal region of the caudal fin. At 10dpa intense blood vessel formation was disseminated throught the regenerated tissue is significantly higher compared to control. A fold increase of of 3.6 in length, 2.8 in junction and 2.6 in size (Fig.2) of the dorsal region compared to cleft region and fold increase of 3.5in length, 2.7in junction, 2.3 in size and ventral region with 2.8 in length, 2.7 in size and 2.1 in size of the blood vessel. Blood vessels branching out thicker through linking new connections between neighboring vessels, switching to form anastomasis in the vascular region under hypoxic condition was found to be superior when compared to control (Fig.1). At 5dpa blood vessel formation is well differentiated under control receiving 0.1%DMSO and 1% CoCl_2_ each other (Fig.5). Consistent exposure to hypoxia showed an increase of 2 fold change in blood vasculature (Fig.1), which was significantly reduced when treated with genistein. At 100µM genistein, fold change of 0.3 times in length and 0.4 times in size of the blood vessel at dorsal region, compared to 0.4 and 0.6 times at cleft region, and 0.34 and 0.52 in ventral region respectively, compared to more than 2 fold increase under 1% CoCl_2_ condition suggesting genistein considerably curtailed blood vessel sprouting (Fig.3). Consistent inhibition of blood vessel by genistein perturbed caudal fin regeneration, in a dose-dependent manner (Fig.6) and these results were compared with fish treated with 1µM SU5416. fold changes were observed in length of blood vessel at 0.1 in dorsal, 0.15 at cleft and 0.1 in ventral regions and size of the blood vessel at 0.14 in doral, 0.2 at cleft and 0.16 at ventral regions were found significantly inhibited compared to control. Due to strong inhibition of SU5416, the constant suppression of blood vessel sprouting effectively arrested regeneration (Fig.3). Collectively these findings demonstrate role for HIF-1α through angiogenesis and provides a rational for therapeutic approach to target HIF-1 for activation.

### 3.2. VEGF signaling during caudal fin regeneration.

Hypoxia is known to induce angiogenesis through activation of VEGF and transcription of HIF-1. To determine the impact of hypoxia induced VEGF signaling could play crucial role during regeneration, adult zebrafish caudal fin was amputated and exposed to cobalt chloride. Similar to hypoxia, zebrafish caudal fin exposed to CoCl_2_ led to extensive blood vessel formation and enhanced regeneration. Relatively at low concentration cobalt chloride considerably induced angiogenesis without any significant impairment in vascular networks, suggesting that blood vasculature was more prone to wound healing. Gene expression of several angiogenic factors along VEGF pathway was obtained from regenerated fins of 1% CoCl_2_ treated adult zebrafish at 5 dpa, and quantified using qPCR. Results indicated an up-regulation of more than 2 fold increase in HIF α/VEGF, VEGFR2 expression (Fig.7), 1.5 fold increase in MMP9, MMP2, 1.8 fold increase in co- NRP1a, FGFR2,EGFR, PDGFR2, less than 2 fold increase in pro-angiogenic genes like angpt1,IGF-1, ITGAV,Tie1. Complete reduction in expression of these mRNA’s is observed in presence of 1μM SU 5416, a specific inhibitor of VEGF-R2, implying VEGF signaling could be important during regeneration. Genistein treatment also resulted in decreased expression of VEGF, VEGF-R2, co-receptor neuropilin 1a and downstream factors such as MMP2, MMP9 and FGFR2 (Fig.8). Gene expression from genistein treated regenerated fin at 5 dpa show decrease in fold change of 0.1 at 25 μM (VEGF, VEGF R2, HIF-1α, NRP1A, TBFb1, TIMP 2A), 0.4 at 50 μM (Notch, HER, HEY, IGF-1) and upto 0.8 fold at 100 μM (VEGF, VEGF-R2, FGFR2, DLL4, MMP9, MMP2, β-CATENIN). We estimated the translated expression of VEGF during CoCl_2_ condition in adult zebrafish using immunoblot technique. Protein detected from time course experiment at 3, 5 and 10 dpa shows significant increase in VEGF expression after exposure to 1% CoCl_2_ whereas the treatment with 1μM SU 5416 confirmed the absence of VEGF (Fig.9). These results demonstrate that CoCl_2_ might induce hypoxia, which could upregulate VEGF, which might significantly contribute to early events of regeneration under hypoxic condition. Western blot analysis performed on genistein treatement shows a decrease VEGF expression at 25, 50, 100 μM concentrations respectively under 1% CoCl_2_ condition (Fig.9) compared to complete inhibition observed with 1μM SU 5416. These results provide convincing evidence that cobalt chloride induced blood vessel sprouting and migration might be mediated by HIF1α- VEGF signaling through activation of HIF-1 pathway.

**Figure.7.**
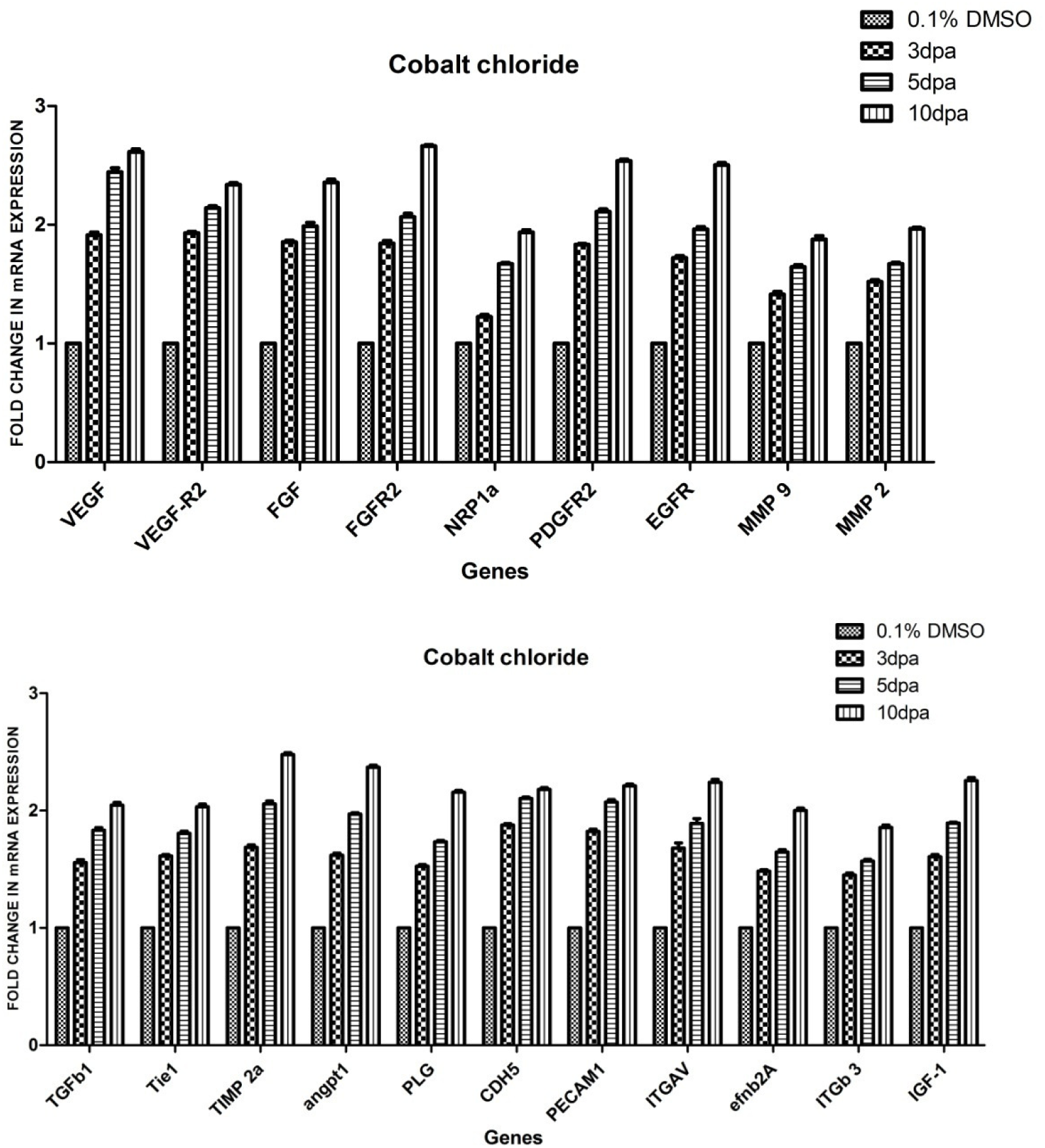
mRNA expression analysis at different time course: VEGF and associated angiogenic factors increased under hypoxic condition (1% CoCl_2_) at 3dpa, 5dpa and 10dpa. at day 3 and day 5 Showing 1.5- 2 fold change increase in of TIE, TIMP2A, PECAM1 IGF-1, TBFb1, and 2-3 fold change increase is observed on day 10.

**Figure.8.**
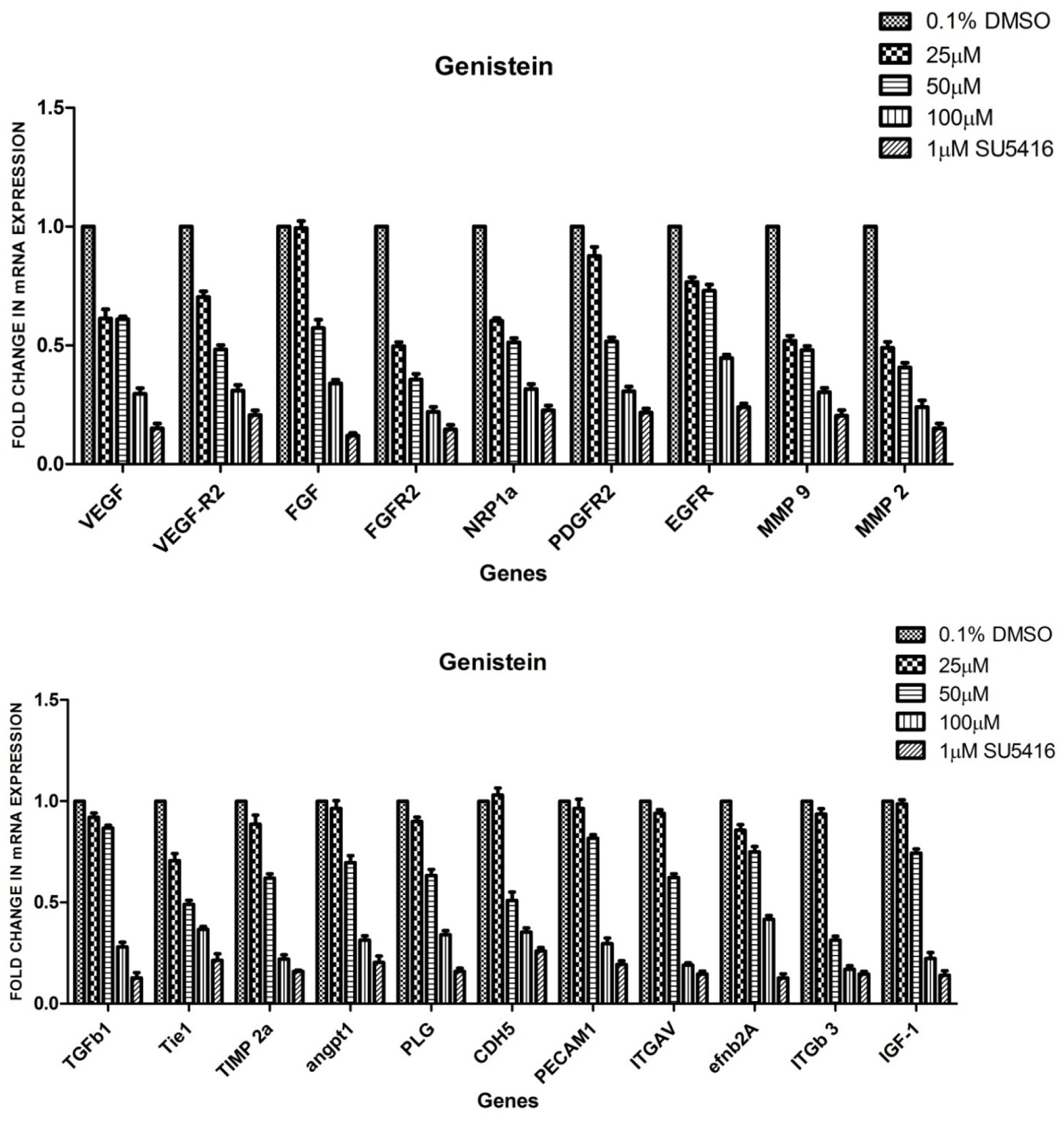
Gene expression analysis of various angiogenic factors under Genistein treatment: The expression is reduced on increasing concentration of genistein on day 5 expressing > 1 at 25µM, at 50µM by 0.5 to 0.9and 100 µM > 0.5 under hypoxic condition (1% CoCl_2_). Genes were expressed upto 0.2 change in 1 µM SU 5416; β-actin was used as housekeeping gene control.

**Figure.9.**
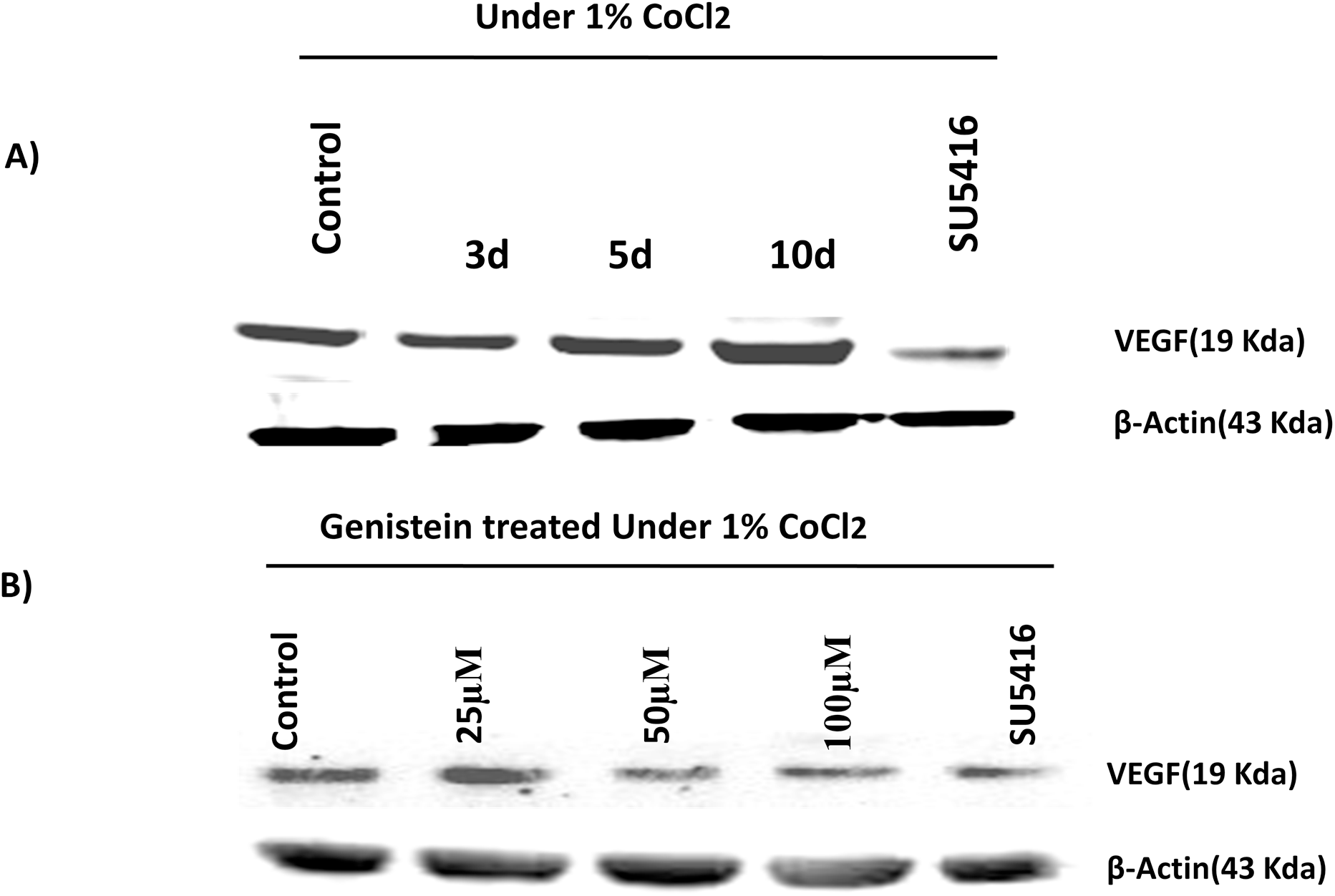
Western blot analysis showing VEGF expression of the regenerated tissue (A) Under hypoxic condition (1% CoCl_2_) at 3 dpa, 5 dpa and 10 dpa (B) Under different concentration of genistein under hypoxic condition. SU 5416 was used as positive control for VEGF; β-actin was used as internal loading control.

## 4. Discussion

Here we have demonstrated caudal fin amputation in adult zebrafish leads to vigorous blood vessel sprouting through CoCl_2_ which enhanced regeneration. Report on hypoxia induced myocardial regeneration in adult zebrafish was positively signed (Jopling C et al., 2012), however in mammals these regenerative events are limited suggest that inhibited or missing factors are of hypoxic response. Similar to hypoxia, CoCl_2_ (1%) led to extensive blood vessel formation, stimulated caudal fin regeneration in adult zebrafish was reported (Mélanie Eyries et al., 2008). Hence these studies on zebrafish present a view point to study wound healing and tissue repair during regeneration. Moreover our data highlighted that inhibitor can effectively block regeneration in zebrafish correlated with hypoxia associated diseases like cancer and diabetic retinopathy. Even though CoCl_2_ plays a beneficial role on immediate treatment to wound healing, prolonged condition might favor cell necrosis. Relatively at low concentration cobalt chloride considerably induced angiogenesis without any significant impairment in vascular networks, suggesting it more prone to wound healing.

Cobalt chloride mimics hypoxic microenvironment, both *in vivo* and *in vitro* (Badr et al., 1999, Wang and Semenza, 1993), via Hif 1α mechanism reported in adult zebrafish (Mélanie Eyries et al., 2008). Cobalt chloride has more affinity towards iron binding site of enzyme blocks hydroxylation activity and prevents binding of HIF-1α- pVHL making HIF-1α expression more stable (Epstein et al. 2001) (Yuan et al., 2003). HIF-1α protein associated with VEGF could be increased by CoCl_2_ induction (Agani et al., 1998). VEGF is upregulated under hypoxic condition via hypoxia inducible factor (Elson et al., 2000; Mace et al., 2007; Pugh CW and Ratcliffe PJ, 2003, Levy AP et al., 1996). Physiological and pathological response of the cells undergoing deprived oxygen supply activates HIF-1α mechanism to maintain vascular homeostasis through VEGF signaling for endothelial cells proliferation thereby promoting wound healing and angiogenesis in regenerated tissue. The data suggest the involvement of VEGF in caudal fin regeneration. Cobalt chloride stimulates angiogenesis by preventing HIF-1α degradation, activating similar set of angiogenic genes regulated under hypoxic conditions (Lee et al., 2001; Vengellur et al., 2003). Molecular insight underlying regeneration involves angiogenic process regulated by various transcriptional and growth factors like VEGF, TGF-β, and PDGF (Galkowska et al., 2006). VEGF is distinctive from other growth factors for its versatile deeds on numerous sequential events, controlling vessel formation during embryonic development (Stojadinovic et al., 2007) sustaining vascular balance in adult organisms by attracting endothelial proliferation and survival (Philip Bao et al., 2009; Susanne Jung and Johannes Kleinheinz., 2013; Glassford et al., 2007), during the course of wound healing. Endothelial cells generated in regenerative tissue are regulated by VEGF expressed in retinal (Takagi et al., 1996) and neural cells [Stein et al., 1995; Levy et al., 1995). Thus amplified VEGF expression regulated by hypoxia might play vital role during tissue regeneration. Previous studies have shown VEGF upregulation under hypoxia in zebrafish larvae (Rathinasamy et al., 2015) is now elucidated in adult Zebrafish caudal fin regeneration.

Generally hypoxia contributes enhanced blood vasculature by altering angiogenic cascade. CoCl_2_ condition boosted endothelial sprouts during regeneration, providing clear transparency by increasing VEGF expression at 3,5,10 dpa respectively. With supporting evidence on fold change increase in blood vessel formation and gene expression indicated cobalt chloride might play pivotal during tissue repair and wound healing present a clinical significance on patients suffering from lack of blood vessel supply to balance blood vessel homeostasis. The feasible mechanism under hypoxia accelerates interface between blood vessel and endothelial cells. Blood vasculature at wounded site is formed with remedial source of target, however sprouting of blood vessel occurred at very early stage of wound healing mechanism. Insufficient blood vessel supply can leave the wound region impaired, and thus early nature of blood vessel formation is essential for its cure. A clinical early induce of blood vessel formation relates the stop point of wound healing cascade involving different steps of blood vessel sprouting and proliferation, which niches the therapy point at damaged site. Although regeneration has proved its impact on gene therapy and tissue engineering studies, which fit to establish major bridge between clinician and research, but understanding its mechanism is a fundamental problem in biology.

Previous report suggests genistein affect receptor tyrosine kinase binding VEFG-R2 to down regulate VEGF in rat (Xu JX et al., 2011), mouse (Farina HG et al., 2006), zebrafish embryo (Bakkiyanathan et al., 2010). This effect was partially abolished in regenerative angiogenesis of adult zebrafish studied under normoxic condition (Rathinasamy et al., 2014). To further explore the importance of VEGF signaling, studies have attempted under hypoxic condition. Investigation on genistein treated zebrafish reduced fin length at 5dpa and down regulated VEGF expression under hypoxia at various concentrations. Corresponding reduction of VEGF under genistein treatment over hypoxic condition could degrade Hif 1α activity. Inhibitory deeds of genistein against CoCl_2_ induced regeneration compared with SU 5416 a VEGF-R2 inhibitor serving as positive control strikes the significance of VEGF signaling during regeneration

**TABLE.2.** List of growth factors studied

|  |  |  |  |  |
| --- | --- | --- | --- | --- |
| Angiogenesis | Notch | Notch - Transcription Factor | Growth factors | Angio/Tissue Maint protein |
| VEGF | NOTCH 1A | hey | FGF2 | CDH5 |
| VEGF-R2 | NOTCH3A | her6 | FGFR2 | PECAM1 |
| NRP1a | DLL4 | NRARPA | PDGFR2 | ITGAV |
| angpt1 |  |  | EGFR | ITGb 3 |
| efnb2A |  |  | TGFb1 |  |
| EPHB4 |  |  | IGF-1 |  |
| MET |  |  |  |  |
| Angiogenesis Inhibitors | Regeneration | Transcription factor | House Keeping gene | Cell surface protein |
| TIMP 2a | SHH | HIF 1α | b-actin | Tie1 |
| p53 | MMP 9 | b-CATENIN |  |  |
|  | MMP 2 |  |  |  |
|  |  |  |  | Enzyme |
|  |  |  |  | PLG |
|  |  |  |  | Angio regulator |
|  |  |  |  | S1PR1 |

FIGURE: 1

## 5. Conclusion

Here we present basic idea on regulation of VEGF signaling under1% CoCl_2_ during tissue regeneration. Observed transcriptional and translational alteration reflects hypoxic response. Increased VEGF expression under hypoxia is consistent leading to tissue regeneration, still needs additional focus to detail the VEGF mechanism during regeneration process. Our findings are significant in demonstrating the impact of VEGF signaling during the process caudal fin regeneration in adult zebrafish under hypoxic stipulation, creates venue for further analyzing the VEGF associated signaling pathways as key biomarker in the process of understanding the regenerative process.

## 6. Acknowledgement

The authors are extremely thankful to the funding provided by UGC-BSR, UGC-SAP, and DST-FIST.

**Ref 1: No: F.7-115/2007(BSR)**

